# Ribosomal DNA arrays are the most H-DNA rich element in the human genome

**DOI:** 10.1101/2024.07.12.602585

**Authors:** Nikol Chantzi, Michail Patsakis, Akshatha Nayak, Austin Montgomery, Ioannis Mouratidis, Ilias Georgakopoulos-Soares

## Abstract

Repetitive DNA sequences can form non-canonical structures such as H-DNA which is an intramolecular triplex DNA structure. The new Telomere-to-Telomere (T2T) genome assembly for the human genome has eliminated gaps, enabling the examination of highly repetitive regions including centromeric and pericentromeric repeats and ribosomal DNA arrays. This gapless assembly allows for the examination of the distribution of H-DNA sequences in parts of the human genome that were not previously annotated. We find that H-DNA appears once every 30,000 bps in the human genome. Its distribution is highly inhomogeneous with H-DNA motif hotspots being detectable in acrocentric chromosomes. Ribosomal DNA arrays in acrocentric chromosomes are the genomic element with the highest H-DNA enrichment, with 13.22% of total H-DNA motifs being found in ribosomal DNA arrays, representing a 42.65-fold enrichment over what would be expected by chance. Across the acrocentric chromosomes we report that 55.87% of all H-DNA motifs found in these chromosomes are in rDNA array loci. The H-DNA motifs are primarily found in the intergenic spacer regions of the ribosomal DNA arrays, generating repeated clusters. We also discover that binding sites for PRDM9, a protein that regulates the formation of double-strand breaks and determines the meiotic recombination hotspots in humans and most mammals, are over 5-fold enriched for H-DNA motifs. Finally, we provide evidence that our findings are consistent in other non-human great ape genomes. We conclude that ribosomal DNA arrays are the most enriched genomic loci for H-DNA sequences in human and other great ape genomes.

## Introduction

The right-handed double-helix structure of DNA, known as B-DNA, was discovered in 1953. Since then, more than 20 non-canonical DNA secondary structures have been reported, including G-quadruplexes, hairpins, cruciforms and triplexes^1^. Sequences that predispose the DNA to non-canonical conformations are known as non-B DNA motifs. Non-B DNA motifs are highly abundant in the human genome and have been associated with a number of biological functions^2–8^.

Mirror repeats consist of two consecutive copies of the same sequence, with one copy being in reverse orientation, separated by an intervening spacer sequence, which lacks symmetry. AG / TC-rich mirror repeats have been shown to fold in intramolecular triple-stranded DNA, also known as H-DNA, in which a strand of DNA with mirror symmetry folds back to itself ^9–11^. Mirror repeat sequences capable of adopting H-DNA conformations are frequently found in mammalian genomes. In the human genome, these sequences have been estimated to occur approximately once every 50,000 base pairs (bps)^12^. H-DNA has been characterized for its functional roles in different biological processes including in gene regulation and in DNA replication ^13–19^. The formation of H-DNA at *MYC* promoter has been shown to interfere with transcription ^13,20^. Studies have also shown that H-DNA is a mutational hotspot for human diseases including cancer ^2,21–23^. For instance, the expansion of GAA·TTC repeats that can form triplex structures have been linked to the etiology of Friedreich’s ataxia ^24,25^.

The genes that encode 45S ribosomal RNA (rDNA) are arranged in long tandem repeats located on the short arms of the acrocentric chromosomes and transcribed by RNA Polymerase I (RNAPI) ^26,27^. Previous work has indicated the presence of H-DNA motifs in human rDNA arrays ^28^. Studies have also located triplex targeting sites at rDNA loci, which can modulate gene expression ^29^. Other studies have found long non-coding RNAs that emerge at rDNA loci, and which can form intermolecular triplex structures at rDNA sites to control the epigenetic state of rDNA genes ^30,31^. Nevertheless, the systematic study of rDNA arrays and the nearby intergenic loci has been hindered by the lack of sequencing technologies that can resolve these most repetitive genomic sequences of the human genome. In the Genome Reference Consortium’s Human Build 38 approximately 8% of the incomplete genomic regions amounted to 8%^32^, underscoring gaps in our understanding of the genome.

The recent completion of the human genome through the Telomere-to-Telomere (T2T) Consortium ^32^ enables the examination of the distribution and frequency of repeat elements in regions of the genome that were only partially annotated previously, including centromeres, telomeres and rDNA arrays. Among these regions, rDNA arrays were previously not resolved due to their highly repetitive nature. A typical diploid human genome contains an average of 315 rDNA copies, with a standard deviation of 104 copies ^33^, while the CHM13 reference genome identified approximately 400 copies ^32^. In the T2T Consortium, rDNA arrays were described as the most complex region of the CHM13 graph and their determination was a major milestone ^32^. In another study they showed that PRDM9 binding sites can be identified at rDNA and mediate frequent double-stranded breaks during meiosis^34^. Other recent efforts to assemble T2T organismal genomes are also ongoing. For instance, recent studies have completed the generation of T2T assemblies for multiple non-human primate species through which we can gain insights into the diversity, evolution and plasticity of different repeats in the primate lineage ^35,36^.

Here, we examined the distribution of H-DNA in human chromosomes using the recent T2T CHM13 gap-less assembly. We report that rDNA arrays have the highest genome-wide density of H-DNA motifs in the human genome (42.65-fold enrichment over background rates) and are orders of magnitude more H-DNA rich than any other genomic compartment. Across the rDNA array copies, the H-DNA motifs are preferentially positioned in specific sites, primarily at the intergenic spacer regions. We show that this phenomenon is consistent in other great ape genomes. We find that binding sites for PRDM9, a protein that is over-represented at rDNA loci and which is responsible for regulating the formation of double-stranded breaks and determining meiotic recombination hotspots in humans and most mammals, are enriched with H-DNA motifs. We conclude that ribosomal DNA arrays contain the highest concentration of H-DNA motifs in the human genome.

## Results

We analyzed the occurrence of mirror repeats, sequences that are repeated with a center of symmetry on the same strand, throughout the T2T reference human genome. We find a total of 1,145,912 mirror repeats, characterized by arm lengths of at least ten bps and spacer lengths less than 8 bps. We selected the subset of mirror repeats with high AG/CT content and small spacer lengths, which can form Hoogsteen bonds and fold in H-DNA structures^37^. Specifically, we chose the mirror repeats that were AG-rich / CT-rich (more than 90% AG/CT content), and filtered out sequences with high AT content (>=80%), in accordance with literature ^12,38^. Using these criteria, we report a total of 205,192 H-DNA motifs, genome-wide. These constitute 34.68 H-DNA motifs per megabase (mB) or 2,078 H-DNA bps per mB. Previous studies had estimated the presence of an H-DNA motif every 50,000 bps ^12^, however here we report that based on the complete human reference genome, that has resolved sequencing gaps and assembled repetitive sequences, there are approximately 1.7 H-DNA motifs per 50,000 bps or 1 H-DNA motif roughly every 30,000 bps.

We were interested to examine the frequency of mirror repeats and H-DNA as a function of spacer and arm lengths. Both mirror repeats and H-DNA motifs show a preference for short spacer lengths. When comparing the distributions of H-DNA motifs to mirror repeats, we find that H-DNA motifs prefer longer spacer lengths (**Figure 1a**). Additionally, we subdivided mirror repeats and H-DNA as a function of arm length (**Figure 1b**). We observe that as expected, the number of mirror repeats and H-DNA motifs declines precipitously with arm length, but do not find statistically significant differences between the density distributions of the arm lengths in mirror repeats and H-DNA motifs (Kolmogorov–Smirnov test, p-value>0.05). We also find that the longest mirror repeat was an H-DNA with arm length of 797bp and spacer length of 1bp. We conclude that H-DNA motifs are more abundant in the human genome than previously estimated, with a small subset having unusually large arm lengths.

**Figure 1:**
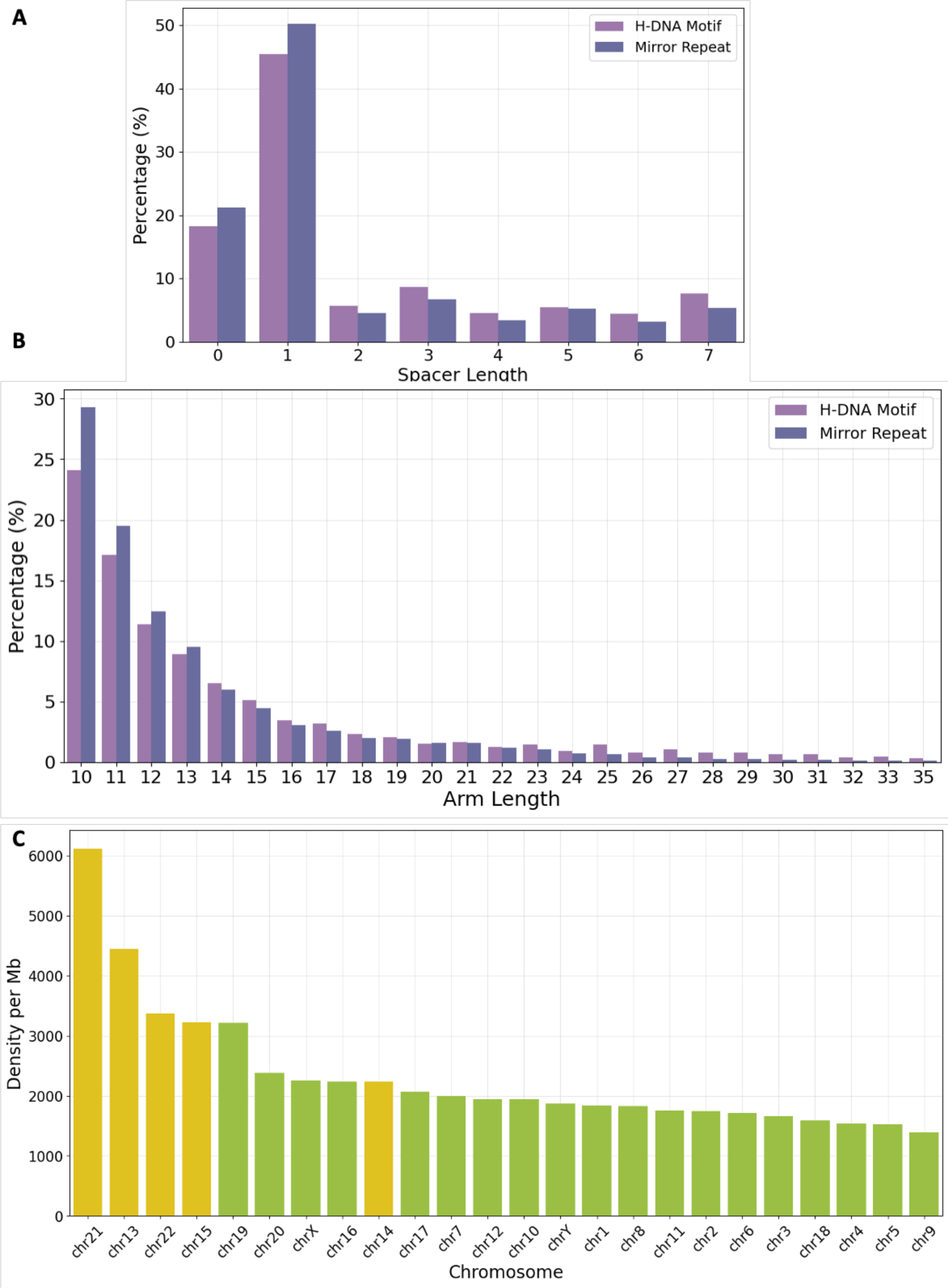
Distribution of mirror repeats and H-DNA motifs in the human genome and across chromosomes. **A.** Distribution density of mirror repeats and H-DNA motifs as a function of spacer length. **B.** Distribution density of mirror repeats and H-DNA motifs as a function of arm length. **C.** Distribution density of H-DNA motif occurrences across human chromosomes. Acrocentric chromosomes with rDNA array loci are colored in yellow.

Next, we were interested in investigating potential differences in the distribution of H-DNA between human chromosomes. We observe marked differences in the density of H-DNA motifs between human chromosomes (**Figure 1c**). We find that four out of the five acrocentric chromosomes showed the highest H-DNA genomic densities, with highest densities among them being observed in chromosomes 21 and 13. This led us to further investigate the reason for the increased frequency of H-DNA sequences in acrocentric chromosomes.

### H-DNA motif frequency is biased between human chromosomes and exhibits hotspots

We investigated if the distribution of H-DNA motifs is uniform throughout the human genome or if it exhibits hotspots and coldspots, loci with either an enrichment or a depletion of H-DNA motifs, respectively. For each human chromosome we generated 2,000 genomic bins of equal length and examined the frequency of H-DNA motifs in each of the bins.

Interestingly, we observe a highly inhomogeneous distribution of H-DNA motifs across the human chromosomes and throughout the genomic bins at individual chromosomes. We find that acrocentric chromosomes, including chromosomes 13, 14, 15, 21 and 22 exhibit localized enrichment peaks for H-DNA sequences, in specific consecutive bins. The highest H-DNA motif enrichments were observed for chromosomes 14 and 22, both showing sharp peaks exceeding 35-fold above background rates. In contrast, chromosomes 15, 21, and 22 had enrichments around 15-fold to 25-fold fold above background rates, but these hotspots covered larger genomic regions of consecutive genomic bins (**Figure 2; Supplementary Figure 1-2**). These enrichments decreased significantly when we examined the distribution of mirror repeats across chromosomes, indicating that the subset of H-DNA motifs are driving the observed enrichment at rDNA arrays (**Figure 2; Supplementary Figure 1-2**).

**Figure 2:**
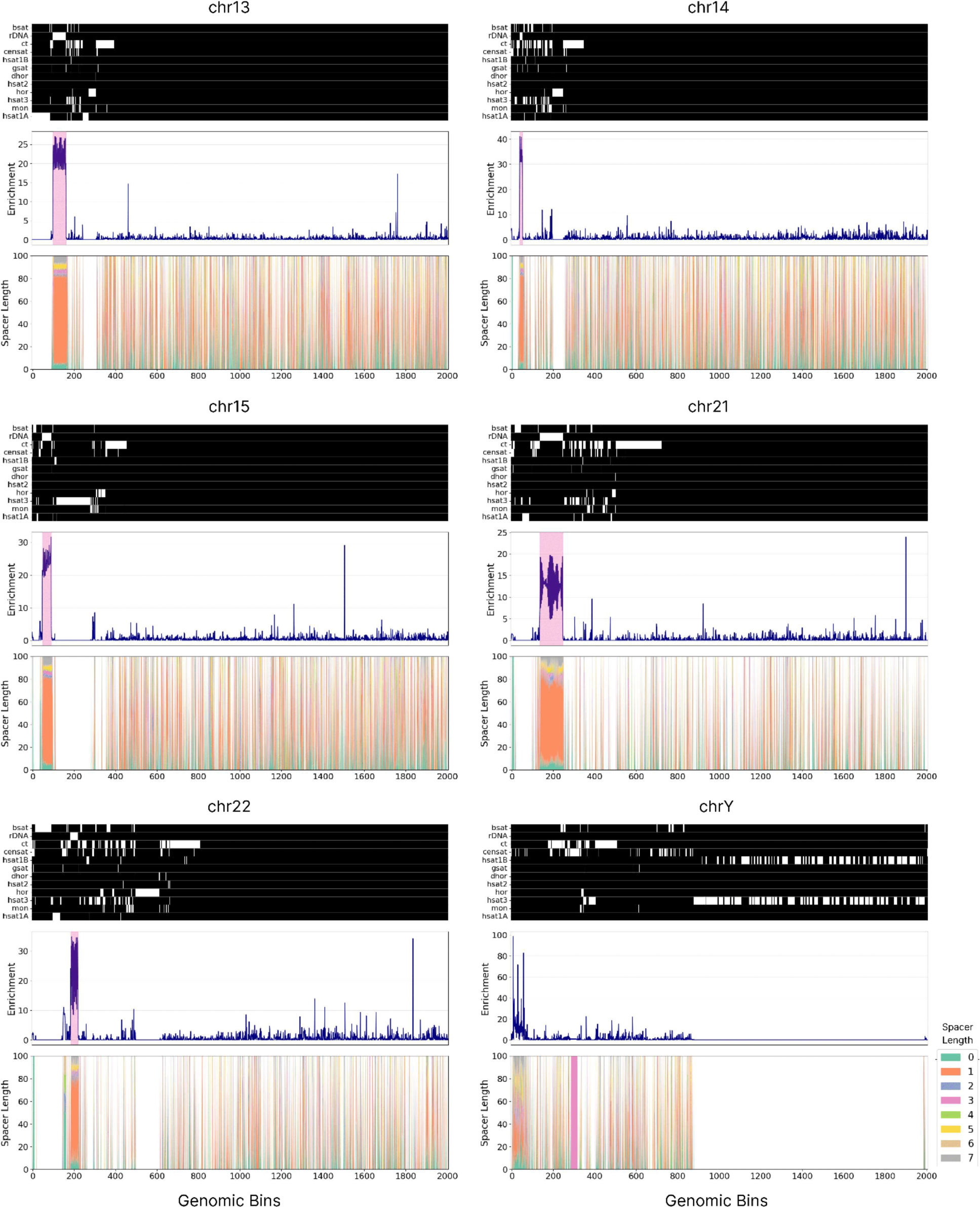
Characterization of H-DNA motifs across chromosomes in the T2T reference human genome. Schematics show the distribution of H-DNA motifs across different human chromosomes. The heatmap shows the different types of pericentromeric and centromeric repeats as well as rDNA arrays, with white color representing presence in that genomic region. Line plots show the H-DNA motif enrichment at each genomic bin for a chromosome. Colored in pink are highlighted the rDNA array loci. Repeats include inactive αSat HOR (hor), divergent αSat HOR (dhor), monomeric αSat (mon), classical human satellite 1A (hsat1A), classical human satellite 1B (hsat1B), classical human satellite 2 (hsat2), classical human satellite 3 (hsat3), beta satellite (bsat), gamma satellite (gsat), other centromeric satellites (censat) and centromeric transition regions (ct).

Across the acrocentric chromosomes 13, 14, 15, 21 and 22, we report that 55.87% of all H-DNA motifs found in these chromosomes are in rDNA array loci. Specifically, in chromosomes 13, and 21, 68% and 71% of H-DNA motifs are found in the rDNA array loci, even though these regions represent a minority of the genomic space of the chromosome. In contrast, the non-acrocentric human chromosomes did not exhibit such pronounced and extended clusters of H-DNA motifs in consecutive genomic bins (**Figure 2; Supplementary Figure 1-2**). These results suggest that H-DNA motifs do not display a uniform distribution throughout the human genome and in individual chromosomes, but instead show highly localized clusters, with most pronounced the clusters at rDNA arrays.

### H-DNA is most enriched at rDNA arrays

Repetitive elements comprise approximately 50% of the human genome ^32^. We were interested to investigate if there are specific genomic elements in which H-DNA motifs form hotspots and compare our findings to the enrichment we observed for rDNA arrays. We aimed to explore the highly repetitive elements recently elucidated by the T2T Consortium. We observed that most H-DNA motif hotspots indeed coincided with rDNA array annotations (**Figure 3**). Even though rDNA array loci represent 0.31% of the human genome, we find that they harbor 13.22% of the total H-DNA motifs. This represents a 42.65-fold enrichment to what would be expected by a random distribution. We compared the density of H-DNA motifs at rDNA loci relative to other genomic sub-compartments. The compared elements included several types of centromeric and pericentromeric repeats, genic regions, enhancers, silencers, rDNA arrays and telomeres. We find that their density is orders of magnitude higher than other sites, with a density of 83.58 bps per kB (**Figure 3**). The second most enriched genomic element from those examined was beta-satellite repeats which showed a 107-fold times lower genomic density than rDNA arrays.

**Figure 3:**
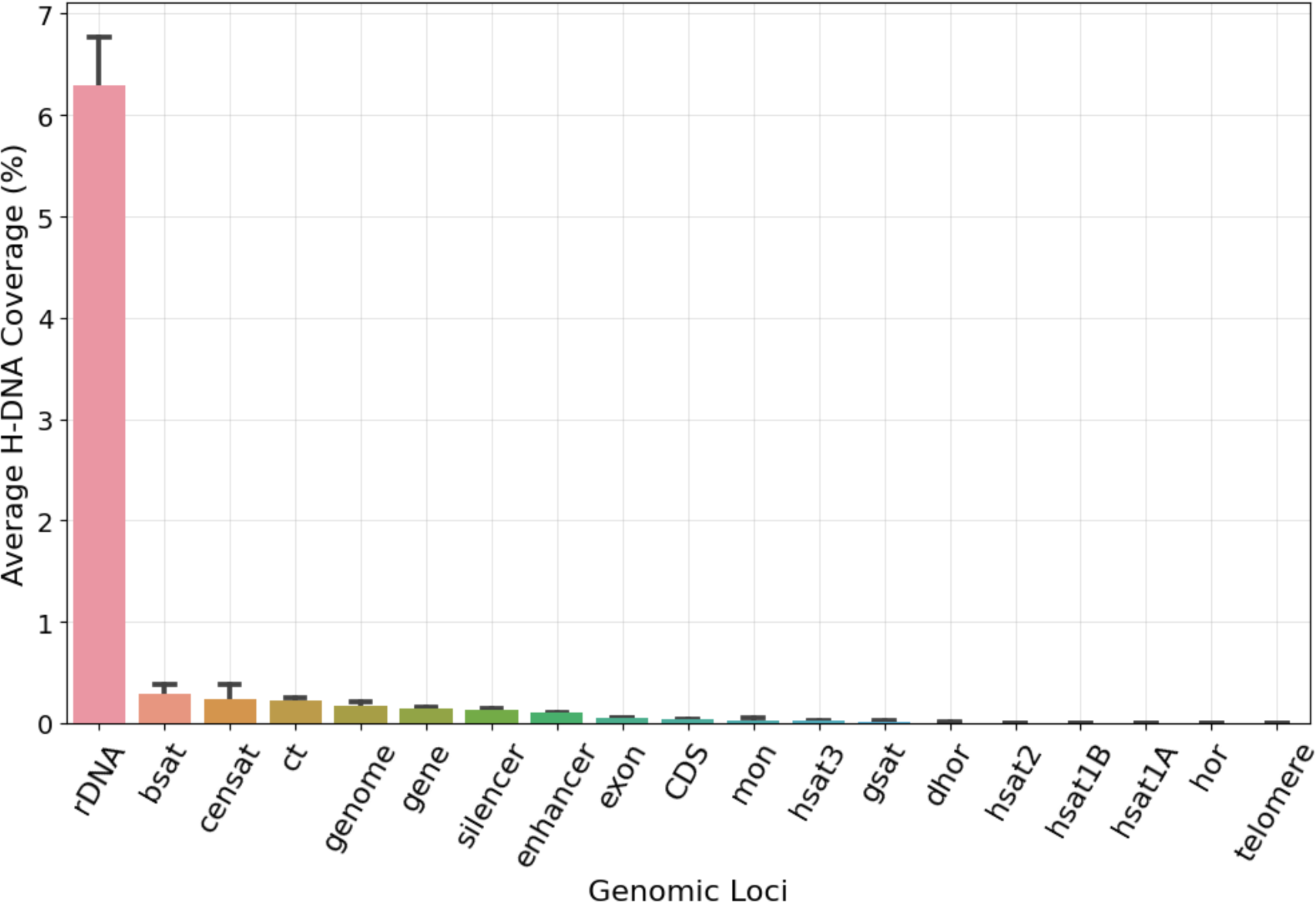
H-DNA motif coverage across human genome sub-compartments and repeats. Repeats include inactive αSat HOR (hor), divergent αSat HOR (dhor), monomeric αSat (mon), classical human satellite 1A (hsat1A), classical human satellite 1B (hsat1B), classical human satellite 2 (hsat2), classical human satellite 3 (hsat3), beta satellite (bsat), gamma satellite (gsat), other centromeric satellites (censat) and centromeric transition regions (ct). Error bars show the 95% confidence interval across instances of each repeat type.

We also examined the frequency of H-DNA motifs across the different types of endogenous repeat elements. Across the endogenous repeat elements, we observe the highest density of H-DNA motifs at retrotransposons, followed by short interspersed nuclear elements (SINEs) (**Supplementary Figure 3a**). We further subdivided the endogenous repeat elements in their sub-families. We find that SINE VNTR Alu (SVA) repeats have the highest H-DNA motif density, followed by Alu repeats (**Supplementary Figure 3b**). These findings are particularly interesting since SVA retrotransposons and Alu repeats are evolutionary young and still active in the human genome ^39,40^. Nevertheless, the observed H-DNA densities at endogenous repeats remained significantly lower than the density at rDNA arrays (**Supplementary Figure 3b, Figure 3)**. We conclude that across the genomic elements examined rDNA is the element with the highest H-DNA density.

### Intergenic loci at rDNA arrays are hotspots for H-DNA

The rDNA encompasses the broader region of ribosomal encoding genes which is highly repetitive and consists of three coding regions, 18S, 5.8S, 28S rRNA and intergenic spacers. As a next step in our analysis, we wanted to examine the positioning of H-DNA motifs within the rDNA tandem array. We examined if H-DNA motifs were more likely to occur in the rDNA genic regions or in the intervening intergenic regions. To that end, we partitioned the rDNA tandem array in rRNA genic regions and intergenic regions and examined the H-DNA density in each respective compartment. The H-DNA signal predominantly originated from the intergenic rather than the rRNA coding, genic regions **(Figure 4a-b**). We also examined the positioning of H-DNA motifs by partitioning the broader rDNA array in 3,500 equally sized bins. The highest number of occurrences of H-DNA motifs were observed in the broader Intergenic spacer (IGS) region downstream of the 28S. Notably, upstream of the 18S genic regions H-DNA are depleted, with re-emergence in proximity to the 28S of the previous adjacent coding array (**Figure 4a-b**).

**Figure 4:**
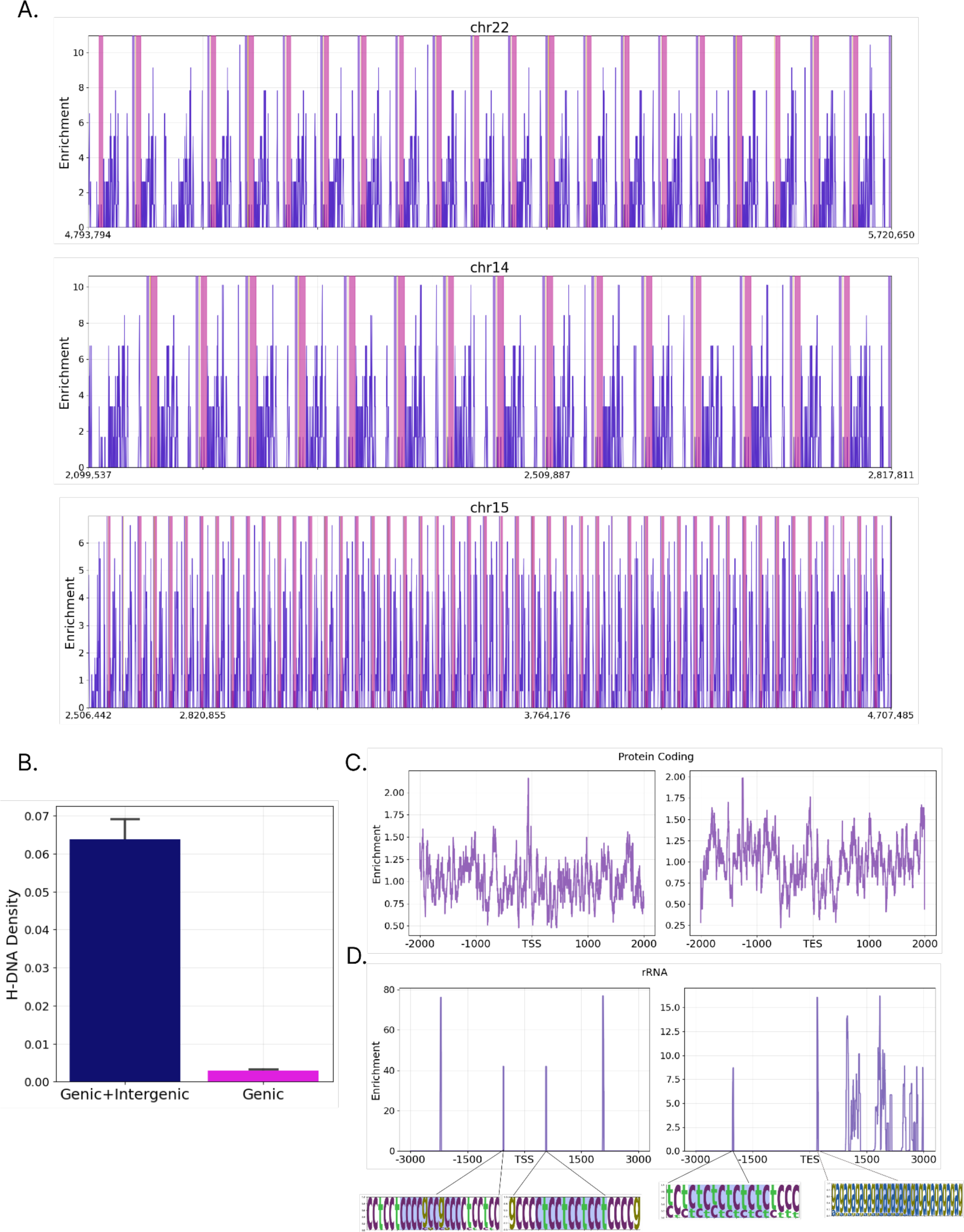
Density of H-DNA in genic and intergenic regions of rDNA arrays. **A.** Enrichment of H-DNA motifs in rDNA arrays in chromosomes 22, 14 and 15. Marked in pink is the location of genes. **B.** Distribution of H-DNA motifs in genic and intergenic regions of rDNA arrays versus in genic regions. **C-D.** H-DNA enrichment relative to the Transcription Start Site (TSS) and the Transcription End Site (TES) of: **C.** protein coding genes and **D.** H-DNA motifs relative to TSS and TES or rDNA arrays. The H-DNA sequences at individual peaks are displayed as positional weight matrices.

Due the high repetitiveness of the rDNA array, the H-DNA motif enrichment phenomenon has a highly predictable periodicity, varying between each chromosome, which is suggestive of the putative regulatory role of H-DNA in rRNA coding genes. We find length 1 being the predominant spacer length of H-DNA motifs in rDNA array loci. Upon examining the extracted H-DNA motifs which emerged in the rDNA array, we noticed that the most frequent H-DNA motif in rDNA arrays, ctctctctgtctgtctctctc, which occurred 1,044 times amongst the examined chromosomes, only appeared 2 times in other parts of the genome (**Supplementary Table 1**).

To further investigate the potential regulatory roles of H-DNA at rRNA coding genes and to obtain a more nuanced representation of the H-DNA motif distribution in relation to the transcription start sites (TSS) and the transcription end sites (TESs), we expanded the respective loci downstream of 28S and upstream of 18S in a window of 4,000bp. The resulting H-DNA distribution verified that most H-DNA motifs are found at 1,000bp downstream of the TES, while the genic area included only a few positions enriched in H-DNA (**Figure 4c**). To investigate if this phenomenon is exclusive to rRNA, we examined the H-DNA distribution of TSS/TES in protein coding genes. The resulting distribution indicated that there is no significant enrichment of H-DNA motifs relative to TSSs and TESs of protein coding genes (**Figure 4c**). These findings indicate a highly biased distribution of H-DNA across the rDNA arrays, with clear preferential positioning, which suggest putative roles of H-DNA motifs in regulating rDNA expression.

### H-DNA is enriched for PRDM9 acting as a recombination hotspot in meiosis

PRDM9 is a zinc finger protein with central roles in determining the location of recombination hotspots across different mammals, including humans ^41,42^. It also mediates programmed DNA double-strand breaks that result in genetic exchange between chromosomes. Previous work has identified the binding motifs using chromatin immunoprecipitation with sequencing experiments ^43^ and shown that PRDM9 binding sites are enriched at rDNA arrays^34^. Additionally, H-DNA is known to be intrinsically mutagenic and cause double-strand breaks ^21,23^.

We hypothesized that PRDM9 binding sites could be enriched at H-DNA motifs because both PRDM9 and H-DNA are preferentially found at rDNA arrays and are linked to double-strand breaks, PRDM9 through its role in meiotic recombination and H-DNA due to its association with increased genomic instability. We examined the genome-wide distribution of H-DNA motifs relative to PRDM9 binding sites and found that H-DNA motifs are over 5-fold enriched at PRDM9 binding sites relative to background rates (**Figure 5a**). We also examined the frequency of PRDM9 binding sites across the rDNA arrays relative to the H-DNA motif loci. We report that 20.6% of PRDM9 binding sites at rDNA loci overlap at least one H-DNA motif and are significantly more likely to overlap H-DNA motifs than expected (Fisher’s exact test, p-value<0.0001; **Figure 5b**). These findings indicate that PRDM9 binding sites are preferentially found in H-DNA motifs. Since H-DNA can cause DNA double strand breaks, this could indicate novel biological roles in mediating meiotic recombination and more work is required to examine if the H-DNA structure formation impacts the genetic exchange between chromosomes during meiosis.

**Figure 5:**
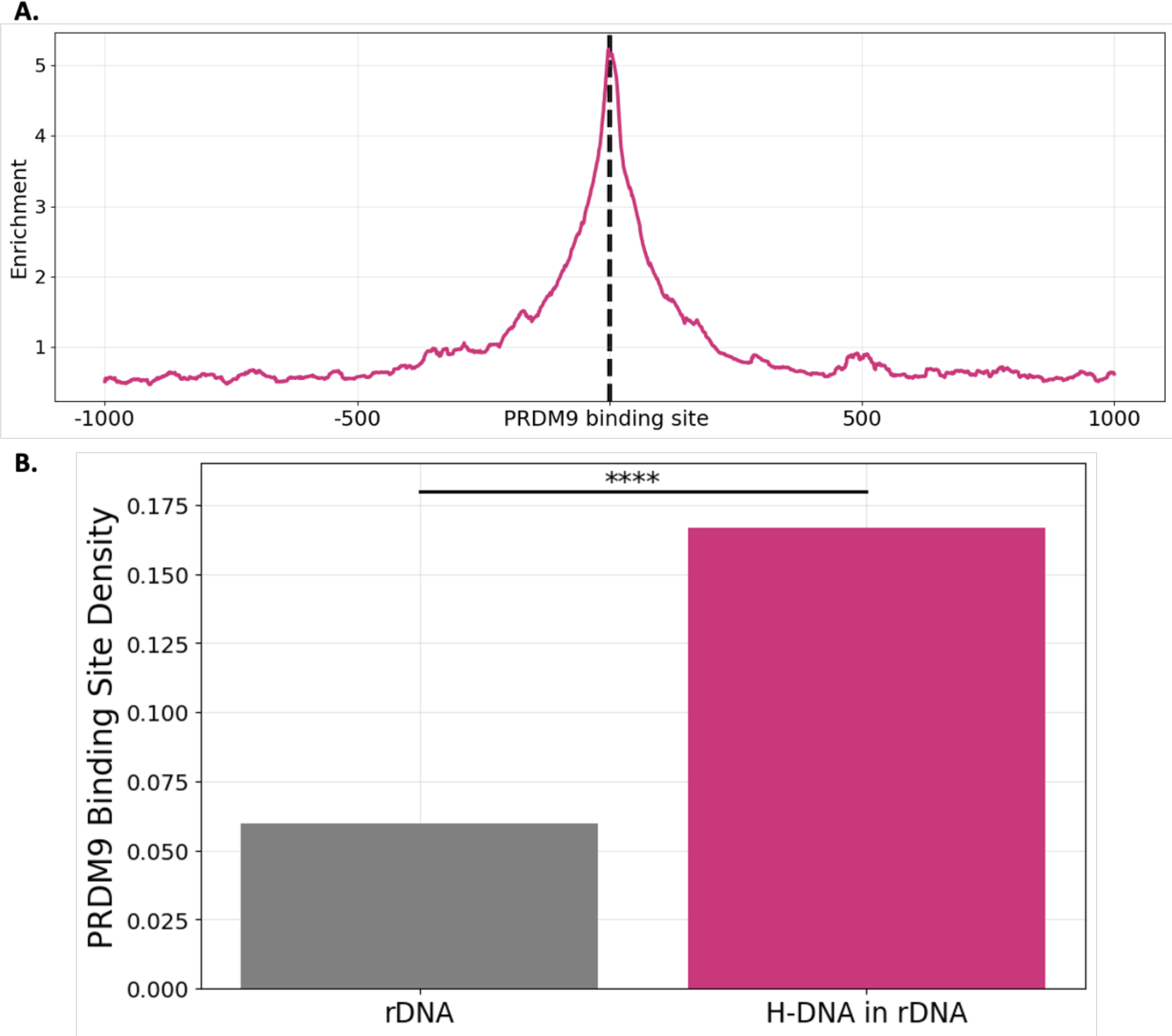
Enrichment of RDM9 motifs at H-DNA loci. **A.** Enrichment of H-DNA motifs relative to the center of PRDM9 binding sites throughout the human genome. **B.** PRDM9 binding site density across the rDNA array and at H-DNA motifs in rDNA arrays.

### H-DNA is highly enriched in rDNA arrays across great ape genomes

Finally, we were interested to investigate if our findings regarding the enrichment of H-DNA in rDNA arrays were also present in other primate species. To provide evidence for the enrichment of H-DNA across great apes, we utilized the T2T primate genomes^35^ recently sequenced and examined the distribution of H-DNA in the genomes of six non-human primates including *Gorilla gorilla* (gorilla), *Pan paniscus* (bonobo), *Pan troglodytes* (chimpanzee), *Pongo abelii* (Sumatran orangutan), *Pongo pygmaeus* (Bornean orangutan) and *Symphalangus syndactylus* (Siamang gibbon).

In these non-human great ape genomes, the number of mirror repeats per genome varies from 3,117,292,070 to 3,545,834,224 in chimpanzee and gorilla species respectively, with a median count of 3,244,508,021.0. For H-DNA motifs, we find that their occurrences per genome ranges between 177,055 and 184,226 in gorilla and Siamang gibbon species respectively, with a median number of H-DNA motifs observed being 178,545.5. Additionally, we find that similarly to humans, for both mirror repeats and H-DNA, repeats with spacer lengths of 0 and 1 bps are the most frequent (**Figure 6a**, **Supplementary Figure 4a**). For both mirror repeats and H-DNA the vast majority of repeats are those with shorter arm lengths and the numbers decline precipitously with increased arm length (**Supplementary Figure 4a**). However, we observe that a subset of H-DNA motifs has extremely large lengths, in several cases exceeding 1,000bps (**Supplementary Table 2**). The largest H-DNA motifs are frequently overlapping microsatellite repeats, which is expected considering the formation of H-DNA secondary structures at such loci, with polypurine/polypyrimidine mirror repeat symmetry ^44^.

**Figure 6:**
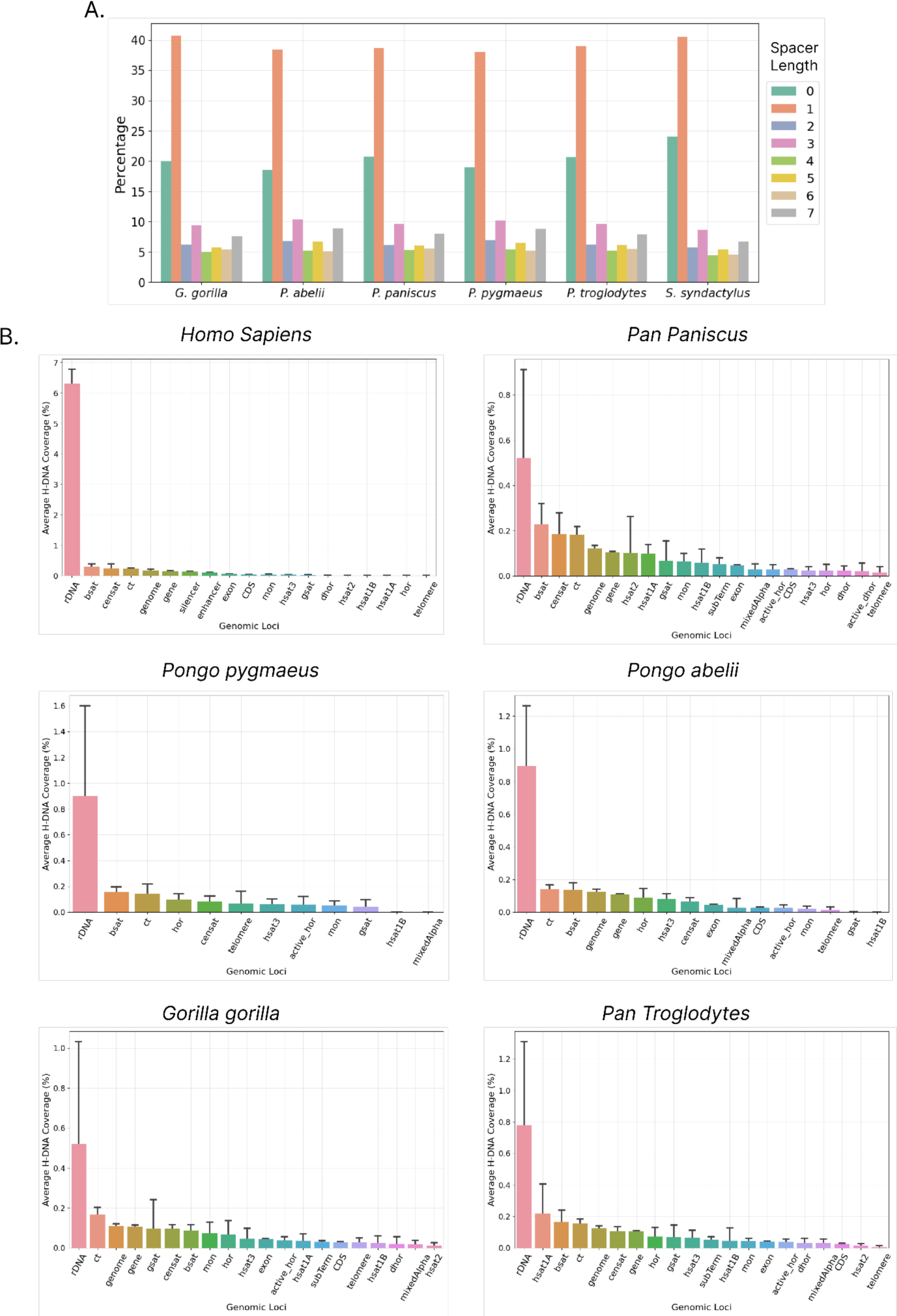
Distribution of H-DNA in Telomere-to-Telomere primate genomes. **A.** Percentage of H-DNA motifs as a function of spacer length. **B.** Repeats include inactive αSat HOR (hor), divergent αSat HOR (dhor), monomeric αSat (mon), classical human satellite 1A (hsat1A), classical human satellite 1B (hsat1B), classical human satellite 2 (hsat2), classical human satellite 3 (hsat3), beta satellite (bsat), gamma satellite (gsat), other centromeric satellites (censat) and centromeric transition regions (ct).

Next, we examined the distribution of H-DNA across different types of genomic compartments and repetitive elements in the great ape genomes. We find consistent results with the analysis performed for the human genome, with H-DNA motifs having the highest density at rDNA loci, across great apes, relative to centromeric and pericentromeric repeats, genic regions and telomeres (**Figure 6b**). However, it should be noted that the genomic annotations of low complexity regions in these genomes may not be as accurately resolved as those in the human genome. Future research is necessary to precisely quantify the enrichment levels of these regions in comparison to other genomic areas. We conclude that rDNA is highly enriched in H-DNA across Great Apes.

## Discussion

Here, we analyzed the T2T reference human genome and found that H-DNA motifs are not homogeneously distributed between and within human chromosomes. We observe that acrocentric chromosomes show some of the highest H-DNA motif enrichments. We provide evidence which indicates that rDNA array loci, at acrocentric chromosomes, are H-DNA motif hotspots, with particularly high density of H-DNA motifs in both human and other great ape primate genomes. When comparing the H-DNA motif density in rDNA arrays to that found in other genomic elements, we observe that rDNA arrays have the highest H-DNA motif density. Interestingly, H-DNA is localized in the intergenic spacer regions between the regions expressing rRNA. We also observe that H-DNA motifs are over 5-fold enriched at PRDM9 binding sites, potentially implicating them in meiotic recombination.

As the ribosome is pivotal in protein assembly, RNA polymerase I accounts for the synthesis of approximately 60% of cellular RNA ^45^. Our discovery of H-DNA motif abundance at rDNA arrays leads us to hypothesize that H-DNA formation has regulatory effects on rRNA expression. Intermolecular triplexes have been shown to form at rDNA loci ^30^. Nevertheless, future work is needed to examine the effects of the formation of H-DNA structures at rDNA loci and decipher their regulatory roles. It will also be important to determine whether H-DNA acts as a regulator of RNA polymerase I and if the positioning of H-DNA motifs impact its activity.

H-DNA is a known mutational hotspot and is associated with genomic instability. Previous studies have shown associations between H-DNA formation and increased mutation rate, as well as links with human diseases ^2,21,23,46–48^. Given the essential role of rDNA in maintaining cellular homeostasis, future research should investigate whether mutations acquired during cancer development or associated with other human diseases at H-DNA sites within rDNA arrays interfere with ribosomal RNA expression.

Furthermore, the enrichment of H-DNA motifs at PRDM9 binding sites, may indicate an involvement of intramolecular triple-stranded DNA structures in meiotic recombination and the exchange of DNA between chromosomes. Previous work has shown that H-DNA causes double-strand breaks *in vivo*^21,23^. Physiological roles of H-DNA formation for the generation of double-strand breaks during meiotic recombination, via PRDM9 binding, is an attractive hypothesis and more work is required to investigate if such mechanisms are modulating meiotic recombination.

Finally, as more T2T genomes are assembled from species of different taxonomic groups, future research will have greater opportunities to uncover the role of H-DNA in influencing eukaryotic evolution and adaptation, particularly as it pertains in the regulation of rDNA expression.

## Methods

### Data Retrieval

We downloaded the reference human genome assembly T2T-CHM13v2.0. Associated files including gene annotation and comprehensive centromere/satellite repeat annotation files were downloaded from https://github.com/marbl/CHM13. The gene annotation GFF file was downloaded from https://ftp.ncbi.nlm.nih.gov/genomes/all/GCF/009/914/755/GCF_009914755.1_T2T-CHM13v2.0/GCF_009914755.1_T2T-CHM13v2.0_genomic.gff.gz and a more comprehensive centromere/satellite repeat annotation file was derived from https://s3-us-west-2.amazonaws.com/human-pangenomics/T2T/CHM13/assemblies/annotation/chm13v2.0_censat_v2.0.bed.

We also downloaded T2T genomes for the following primates ^35,36^, *Gorilla gorilla, Pan paniscus*, *Pan troglodytes*, *Pongo abelii*, *Pongo pygmaeus* and *Symphalangus syndactylus* from https://github.com/marbl/Primates. Associated files including genome annotations were downloaded from https://github.com/marbl/t2t-browser, from which we also derived annotations for centromere/satellite repeats and gene annotations.

PRDM9 binding sites throughout the human genome were derived from https://zenodo.org/records/7692555/files/SupplementaryFile7.chm13v2.PRDM9.tsv?download=1 as estimated in Guarracino et al. ^34^.

### Identification of H-DNA motifs

A mirror repeat was defined as a sequence repeated with a center of symmetry on the same strand with arm sizes of at least 10bp and spacer lengths of less than eight bps. A subset of mirror repeats, known as H-DNA, is predisposed to forming triple helical structures through Hoogsteen bonds. H-DNA sequences were defined with a high AG/CT content exceeding 90%, arm lengths of at least 10bp, spacer sizes of less than 8bp, and excluding sequences with equal or greater than 0.8 total AT content, in accordance with literature ^12,38^. Mirror repeats were identified with the tool non-B-gfa ^38^ and H-DNA motifs were filtered based on the selected parameters. Furthermore, consensus sequences containing N or other characters not originating from the nucleotide alphabet {’A’, ‘G’, ‘C’, ‘T’} were filtered out from the extracted H-DNA dataset. The frequency of mirror repeats and H-DNA motifs with different spacers and arms was calculated.

H-DNA motif density was calculated as the number of H-DNA motif bps over the number of base pairs examined. The H-DNA motif density across genomic subcompartments was assessed by calculating the ratio of the length of H-DNA motif overlaps to the total length of each subcompartment. Subcompartment coordinates were derived from the corresponding GFF files, and any overlapping annotations within a subcompartment were consolidated.

### Generation of genomic bins

For each chromosome for the human T2T genome, we generated N=2,000 genomic bins of equal length. Each H-DNA start and end coordinate was assigned at a particular bin from 1 to 2,000, according to the following formula:

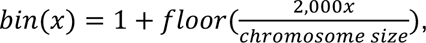

where x is the start or end coordinate of the H-DNA motif. Subsequently, for each bin the total number of H-DNA occurrences was estimated by summing the total distinct start or end coordinates within a particular bin. In cases where the start and the end of the H-DNA were assigned a different bin, the H-DNA was counted in all the intermediate bins. We calculated the number of bps covered by H-DNA motifs, for each H-DNA spacer length.

The wordcloud plots were constructed by keeping only the arm sequences of the H-DNA motifs at the vicinity of the TSS (+/- 2,000bp) and at the vicinity of the TES (+/- 2,000bp) and estimating the most frequent arm sequences across all instances of H-DNA motifs.

We also examined the positioning of H-DNA motifs by partitioning the broader rDNA array in 3,500 equally sized bins. The H-DNA motifs table was filtered in order to contain solely H-DNA motifs belonging to the corresponding rDNA array on each chromosome. The highlighted areas correspond to rRNA genic compartments derived from the respective GFF NCBI annotations. In particular, depending on the position of the genic rRNA compartment in the adjacent rDNA array, different colors were used to indicate 18S, 5.8S, 28S, respectively. Furthermore, we calculated the total number of H-DNA motifs occurring in each bin in the rDNA array and the final enrichment was derived by dividing the total number of observed H-DNA across the spanning rDNA array.

### Estimation of H-DNA density relative to PRDM9 binding sites

To investigate the relationship between H-DNA motifs and PRDM9 binding sites, we generated local windows around the center of the PRDM9 binding sites and measured the distribution of H-DNA bps across the window. Prior to density estimation, the PRDM9 overlapping motifs were merged into a superset interval. The enrichment was calculated as the number of occurrences at a position over the mean number of occurrences across the window.

## Data and Code Availability

The GitHub code and all the related material is provided at: https://github.com/Georgakopoulos-Soares-lab/hdna_rdna_homo_sapiens

## Author Contributions

N.C., and I.G.S. conceived the study. I.G.S. supervised the study and provided the resources. N.C. and I.G.S. generated the figures and tables. N.C., and I.G.S. wrote the manuscript with help from M.P., A.N.,, A.M., and I.M..

## Supplementary Material

**Supplementary Figure 1:**
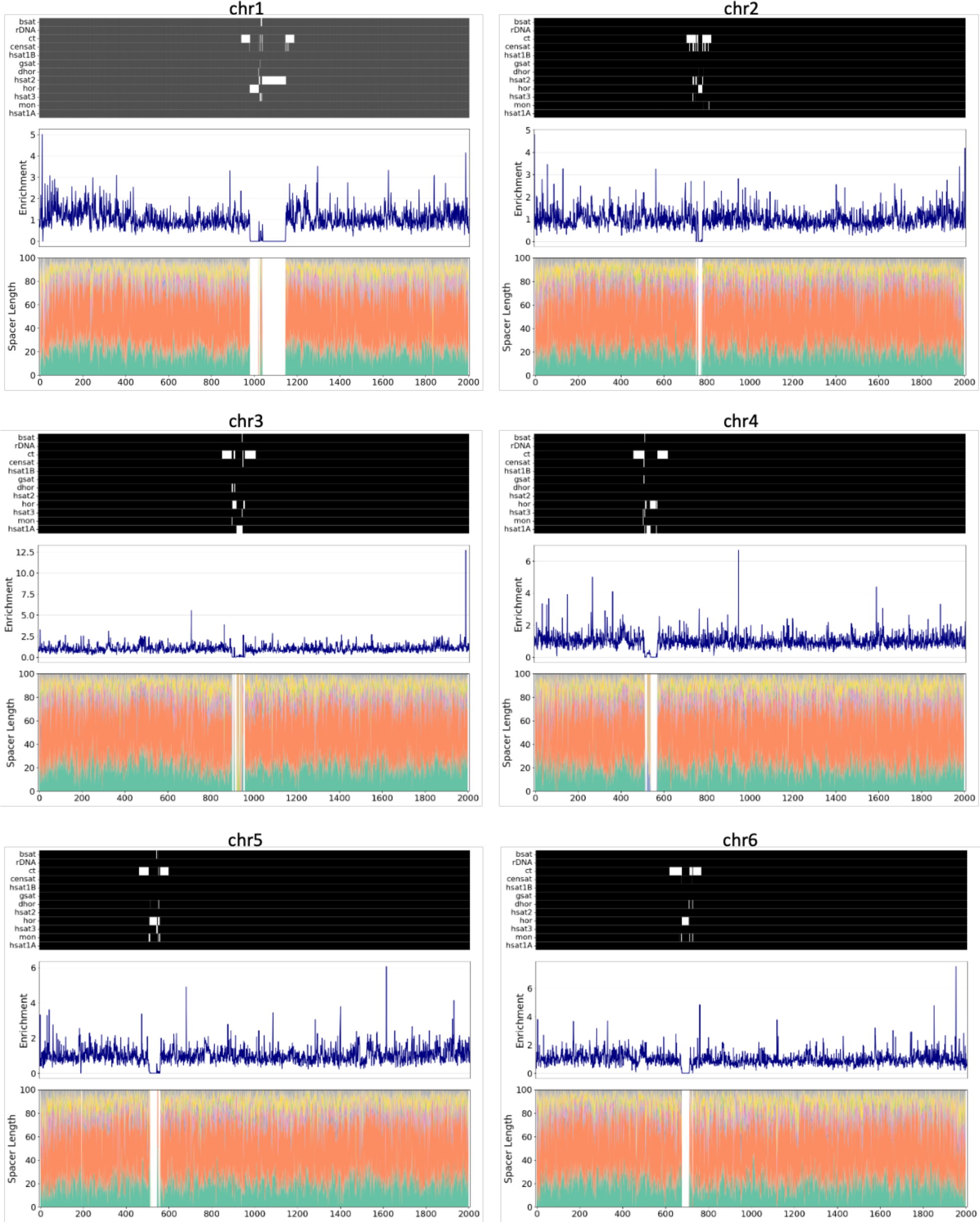
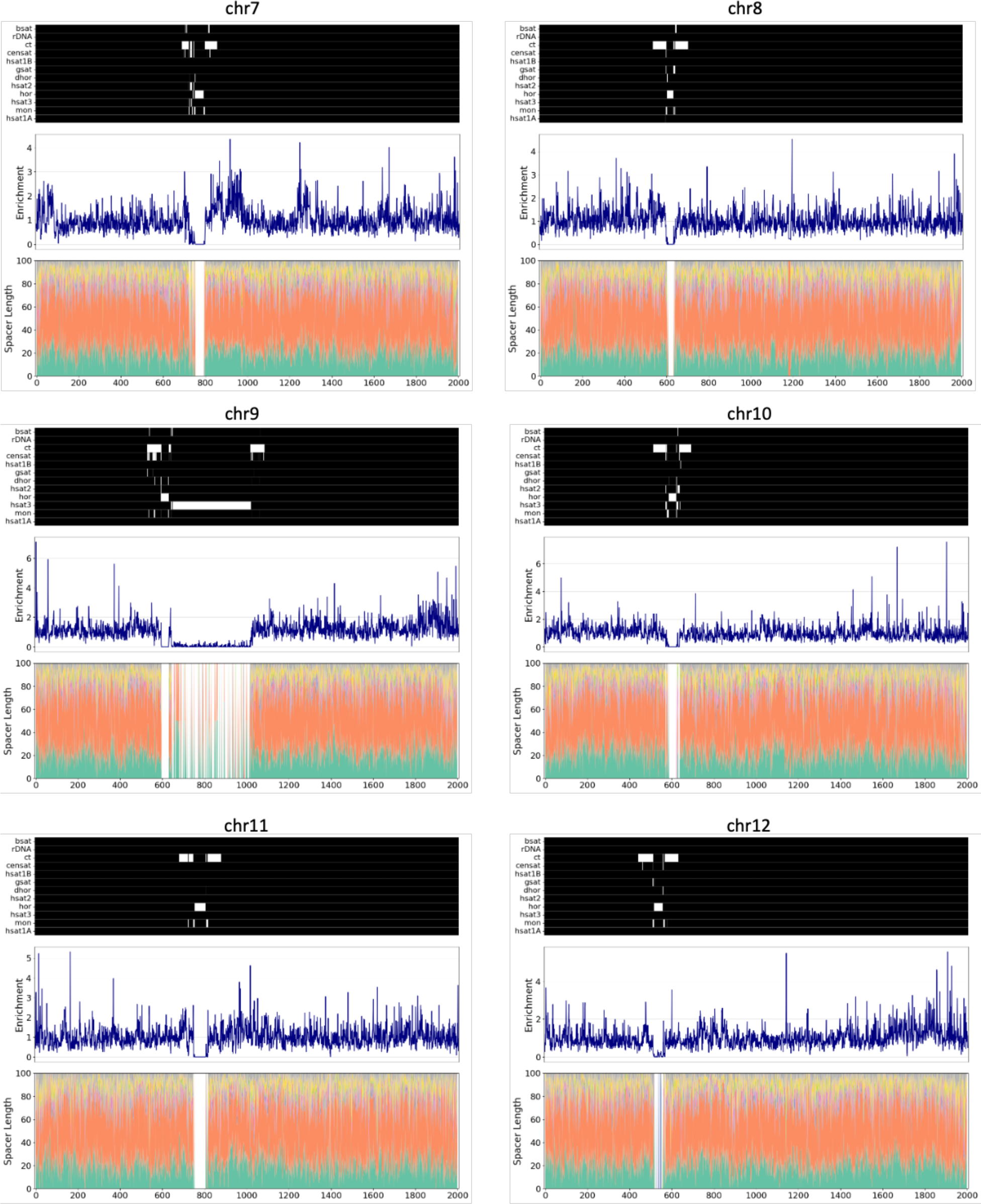
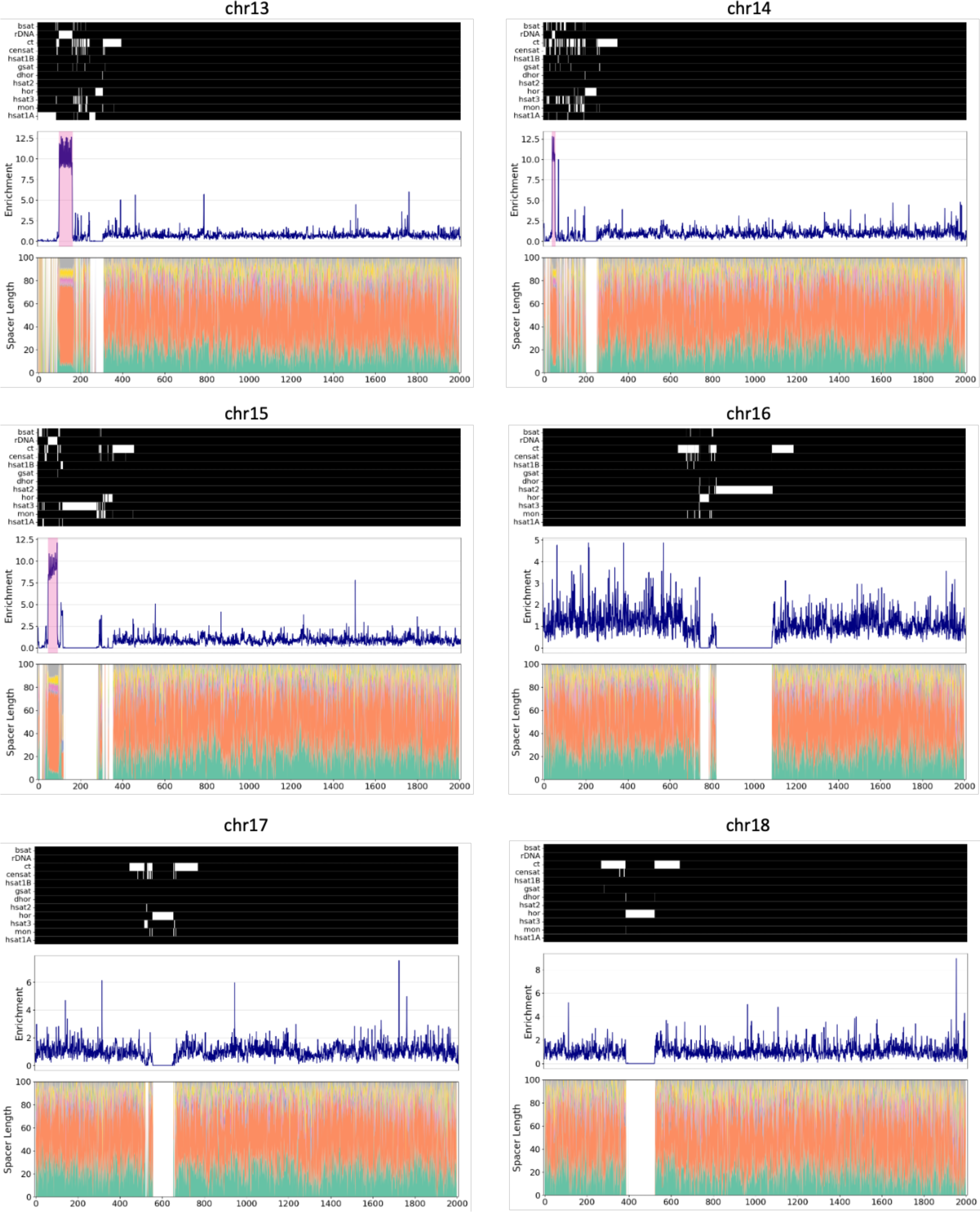
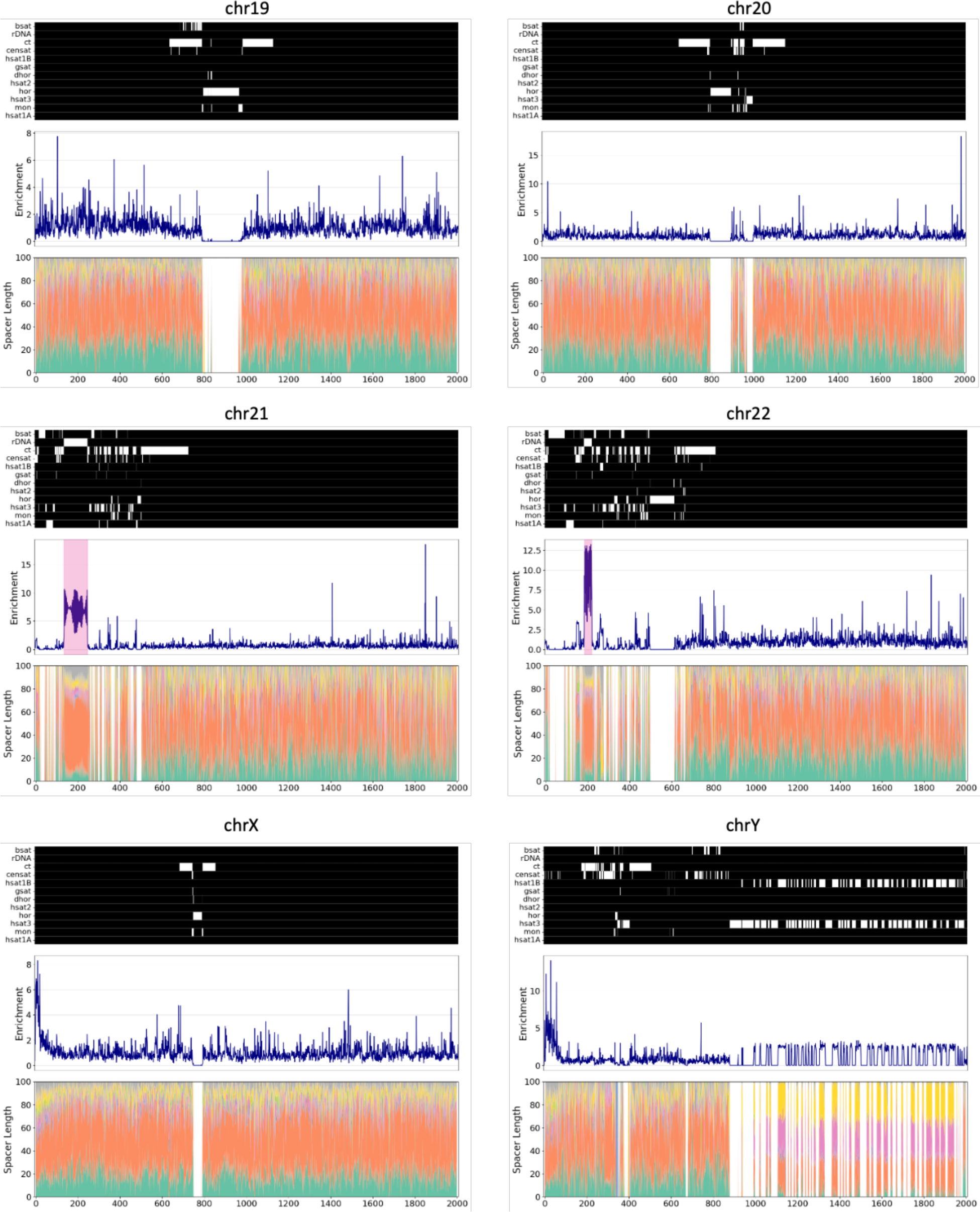
Mirror repeat profile in different human chromosomes. Schematics show the distribution of mirror repeats across different human chromosomes. The heatmap shows the different types of pericentromeric and centromeric repeats as well as rDNA arrays, with white color representing presence in that genomic region. Line plots show the mirror repeat enrichment at each genomic bin for a chromosome. Colored in pink are highlighted the rDNA array loci. Repeats include inactive αSat HOR (hor), divergent αSat HOR (dhor), monomeric αSat (mon), classical human satellite 1A (hsat1A), classical human satellite 1B (hsat1B), classical human satellite 2 (hsat2), classical human satellite 3 (hsat3), beta satellite (bsat), gamma satellite (gsat), other centromeric satellites (censat) and centromeric transition regions (ct).

**Supplementary Figure 2:**
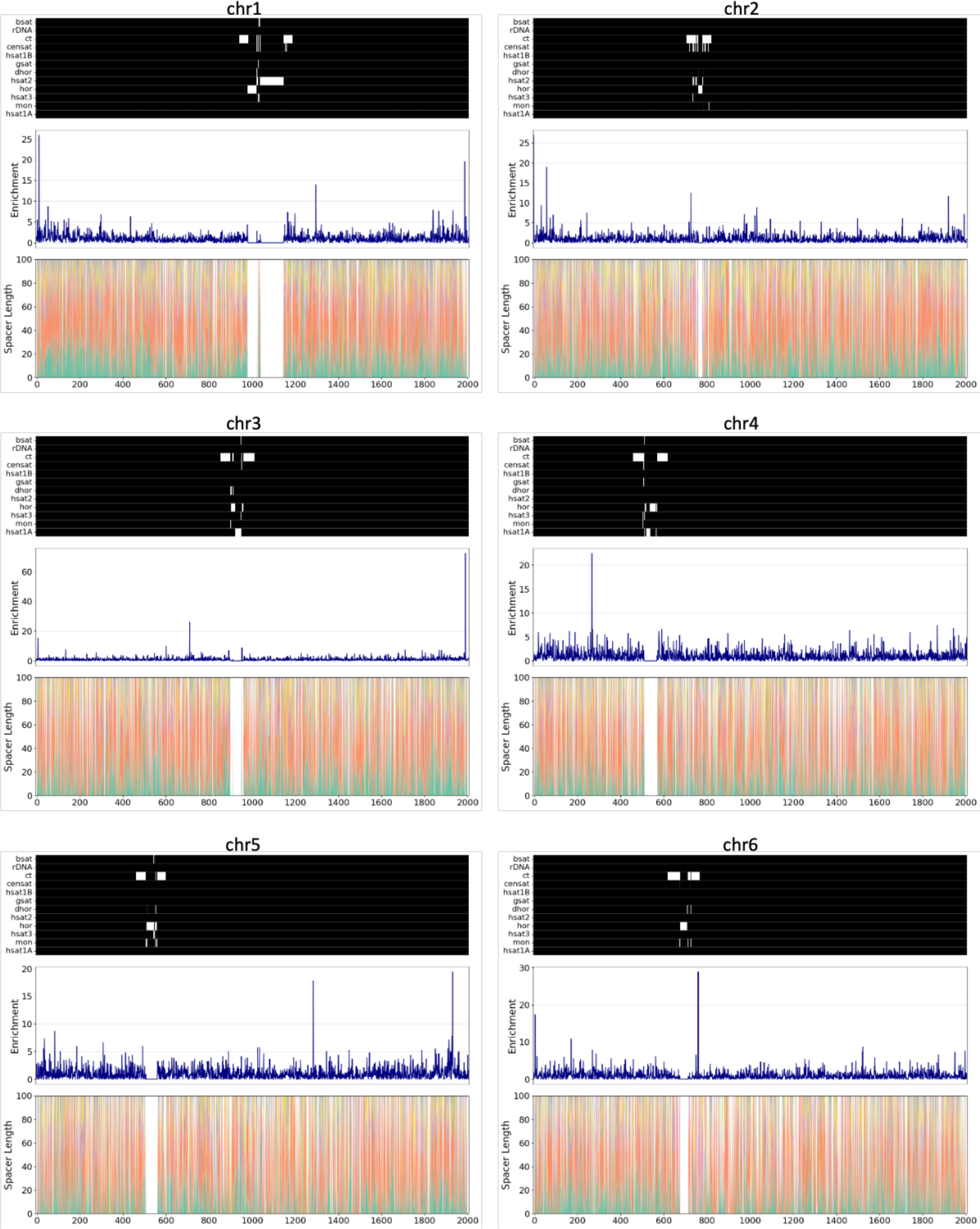
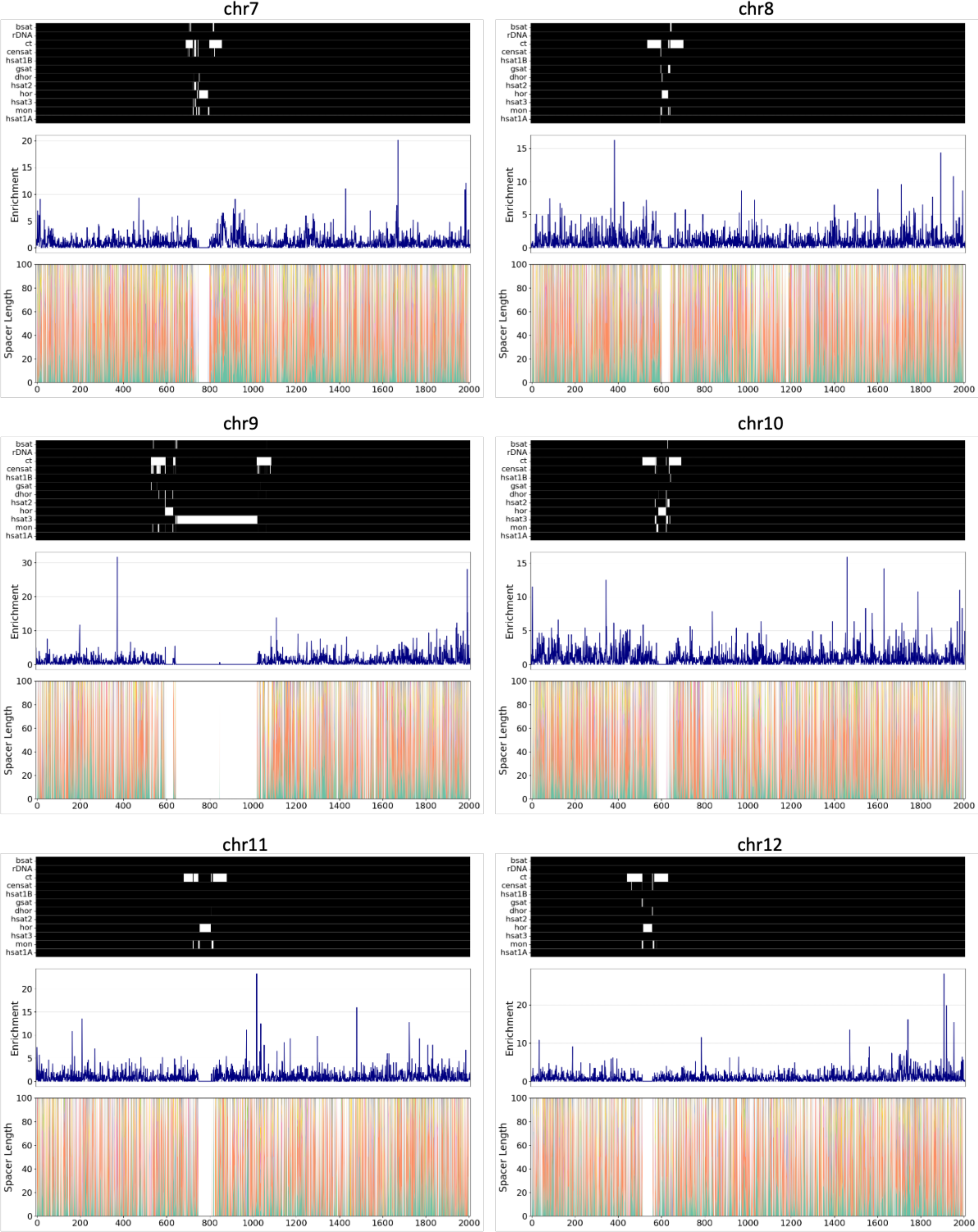
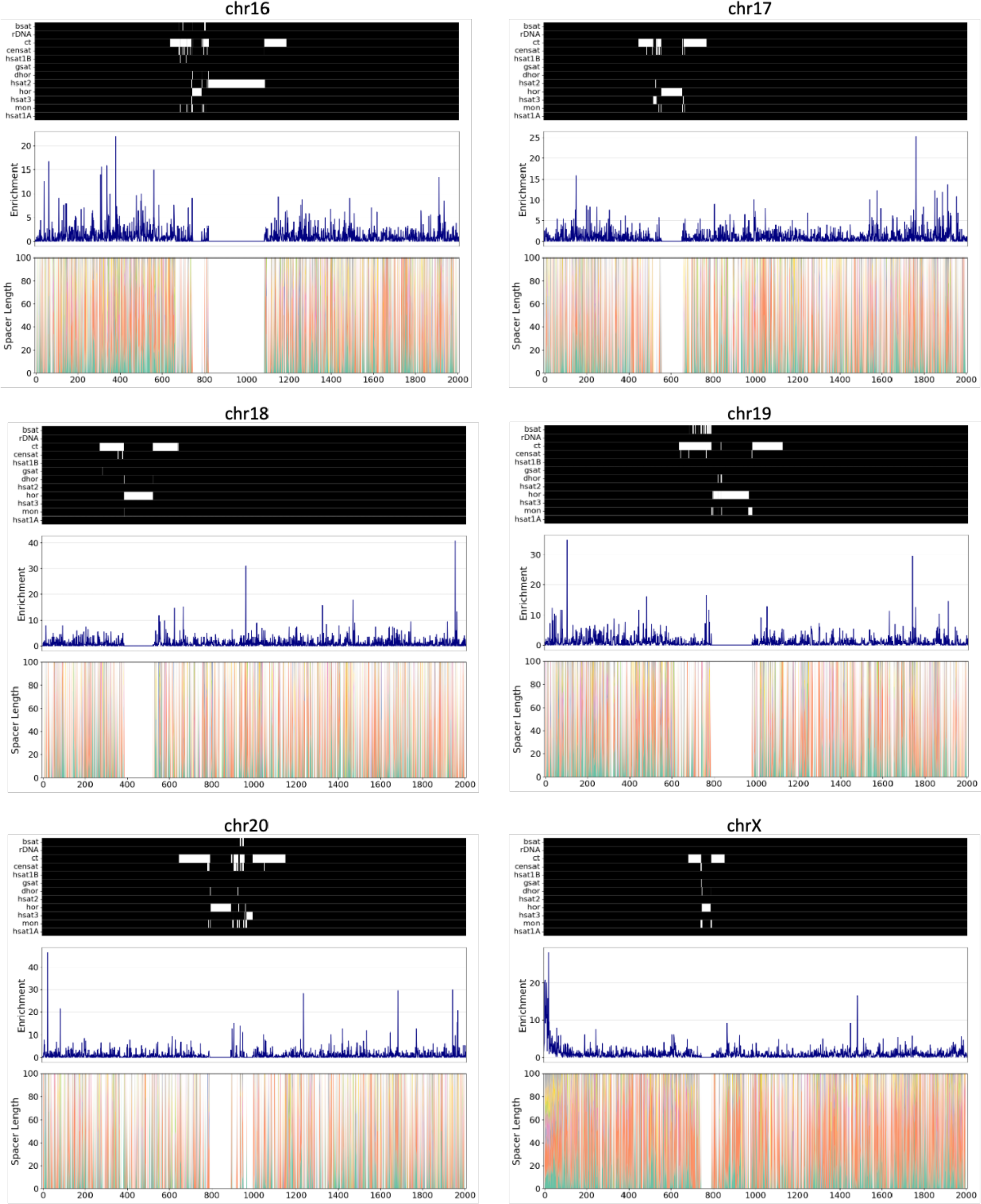
H-DNA profile in different human chromosomes. Schematics show the distribution of H-DNA motifs across different human chromosomes. The heatmap shows the different types of pericentromeric and centromeric repeats as well as rDNA arrays, with white color representing presence in that genomic region. Line plots show the H-DNA motif enrichment at each genomic bin for a chromosome. Colored in pink are highlighted the rDNA array loci. Repeats include inactive αSat HOR (hor), divergent αSat HOR (dhor), monomeric αSat (mon), classical human satellite 1A (hsat1A), classical human satellite 1B (hsat1B), classical human satellite 2 (hsat2), classical human satellite 3 (hsat3), beta satellite (bsat), gamma satellite (gsat), other centromeric satellites (censat) and centromeric transition regions (ct).

**Supplementary Figure 3:**
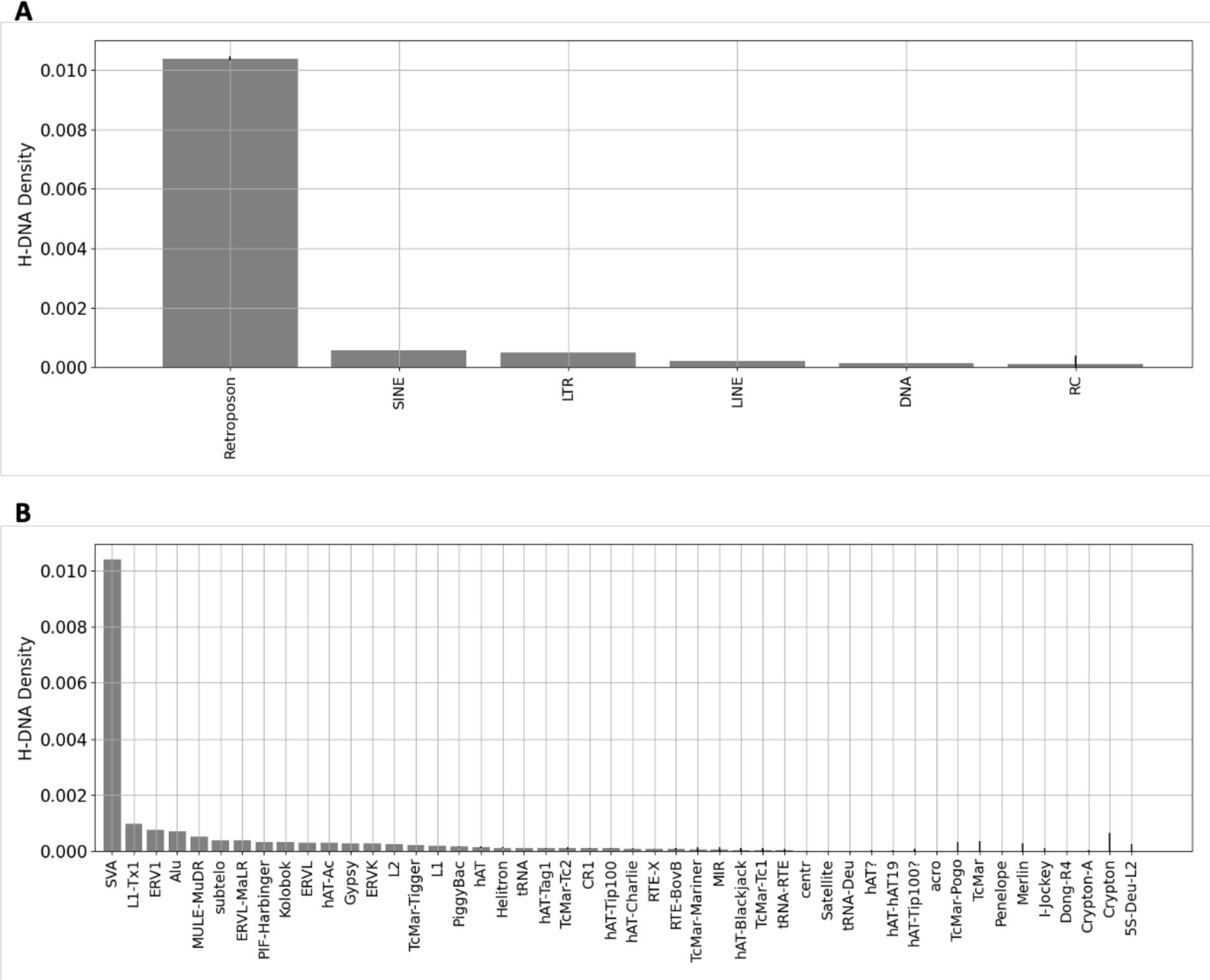
H-DNA Density across transposable elements in the CHM13 reference human genome. **A.** Distribution of G4s in transposable elements and satellite repeats. **B.** Distribution of G4s in transposable elements sub-categories. Error bars represent standard deviation from bootstrapping with replacement (N=1,000).

**Supplementary Figure 4:**
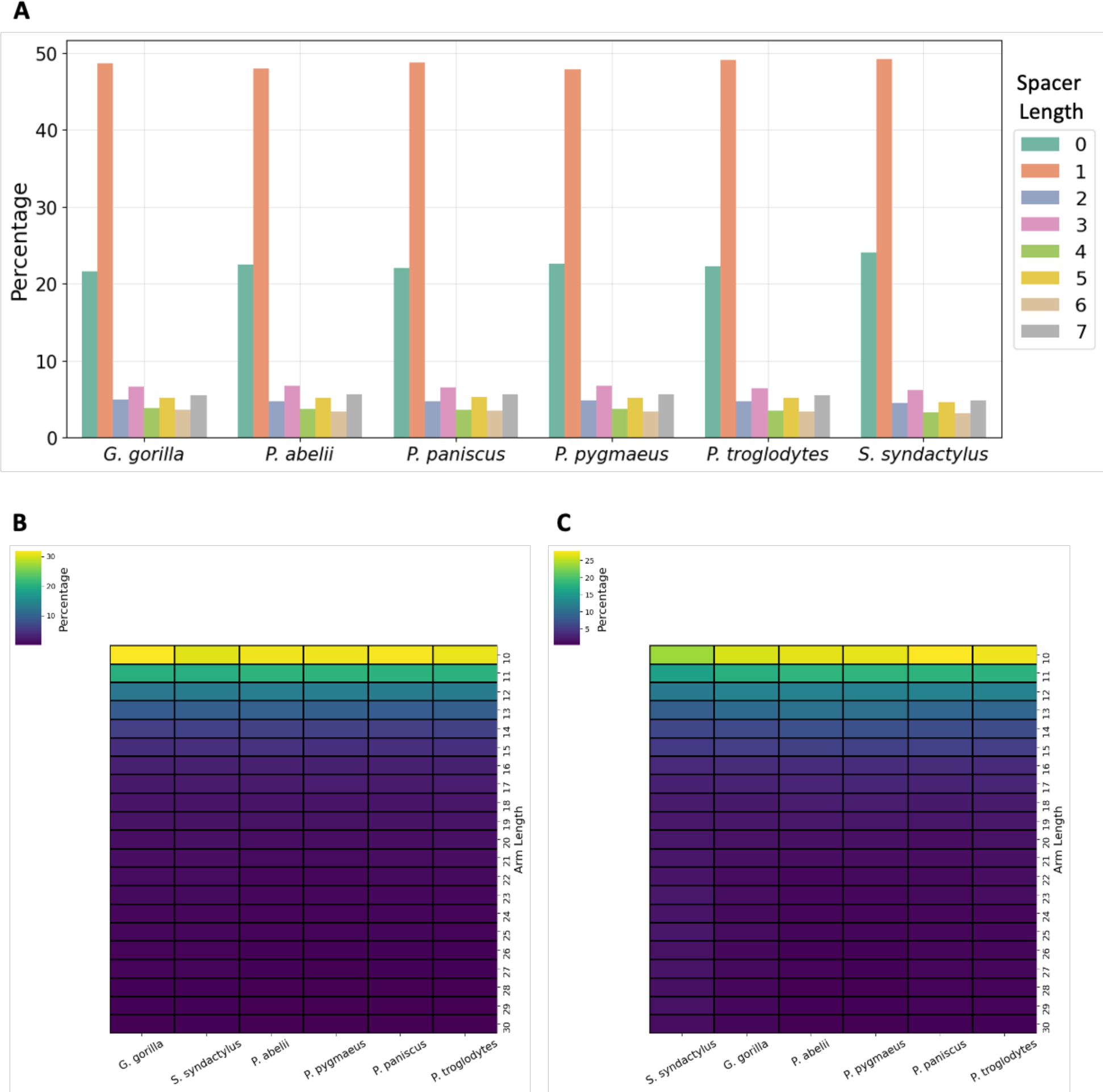
Sequence analysis of mirror repeats and H-DNA motifs. **A.** Percentage of mirror repeats as a function of spacer length. **B.** Percentage of mirror repeats as a function of arm length. **C.** Percentage of H-DNA motifs as a function of arm length.

**Supplementary Table 1:**
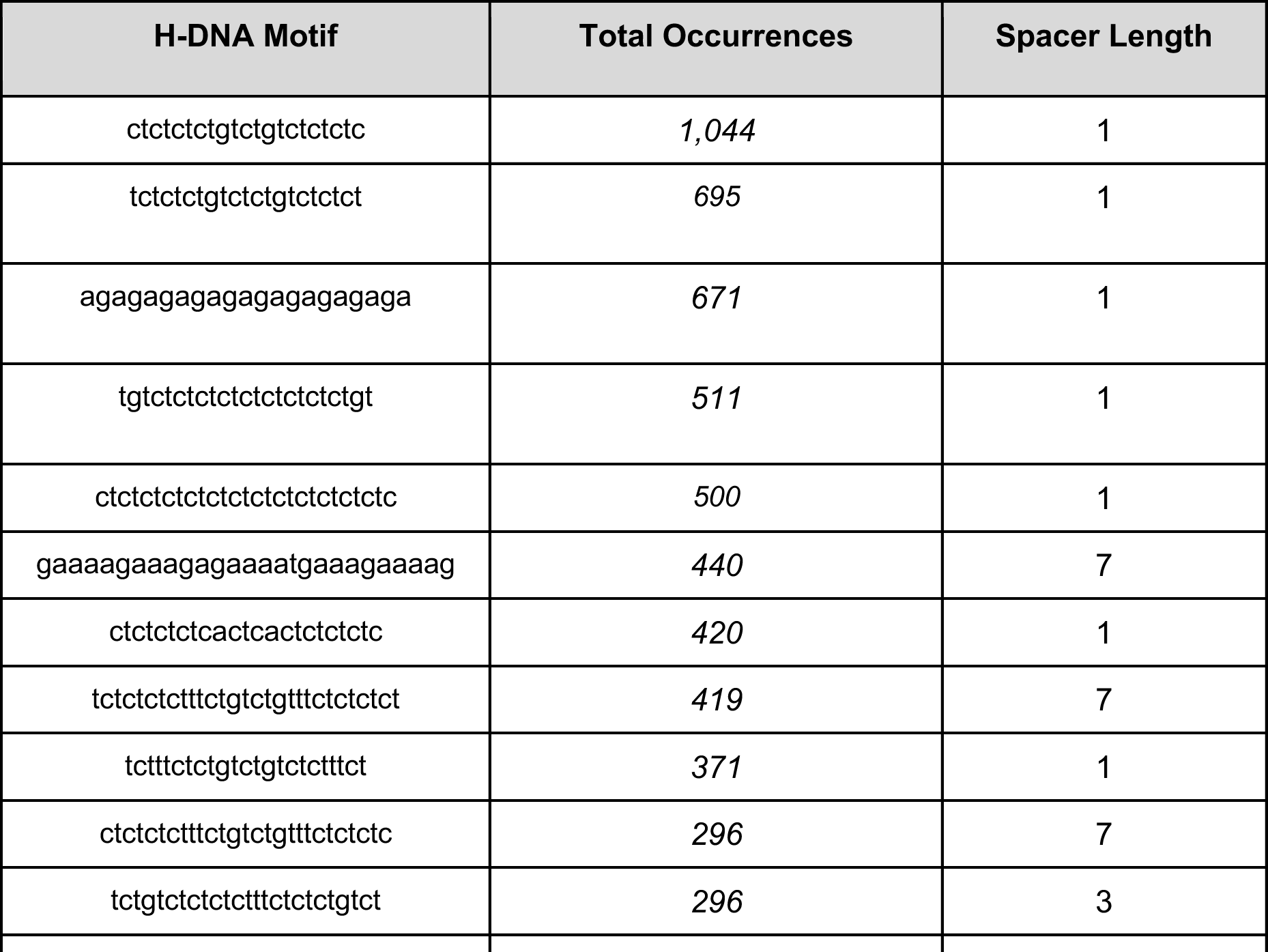
H-DNA motifs with highest number of counts in human rDNA array and the intervening intergenic regions.

**Supplementary Table 2:**
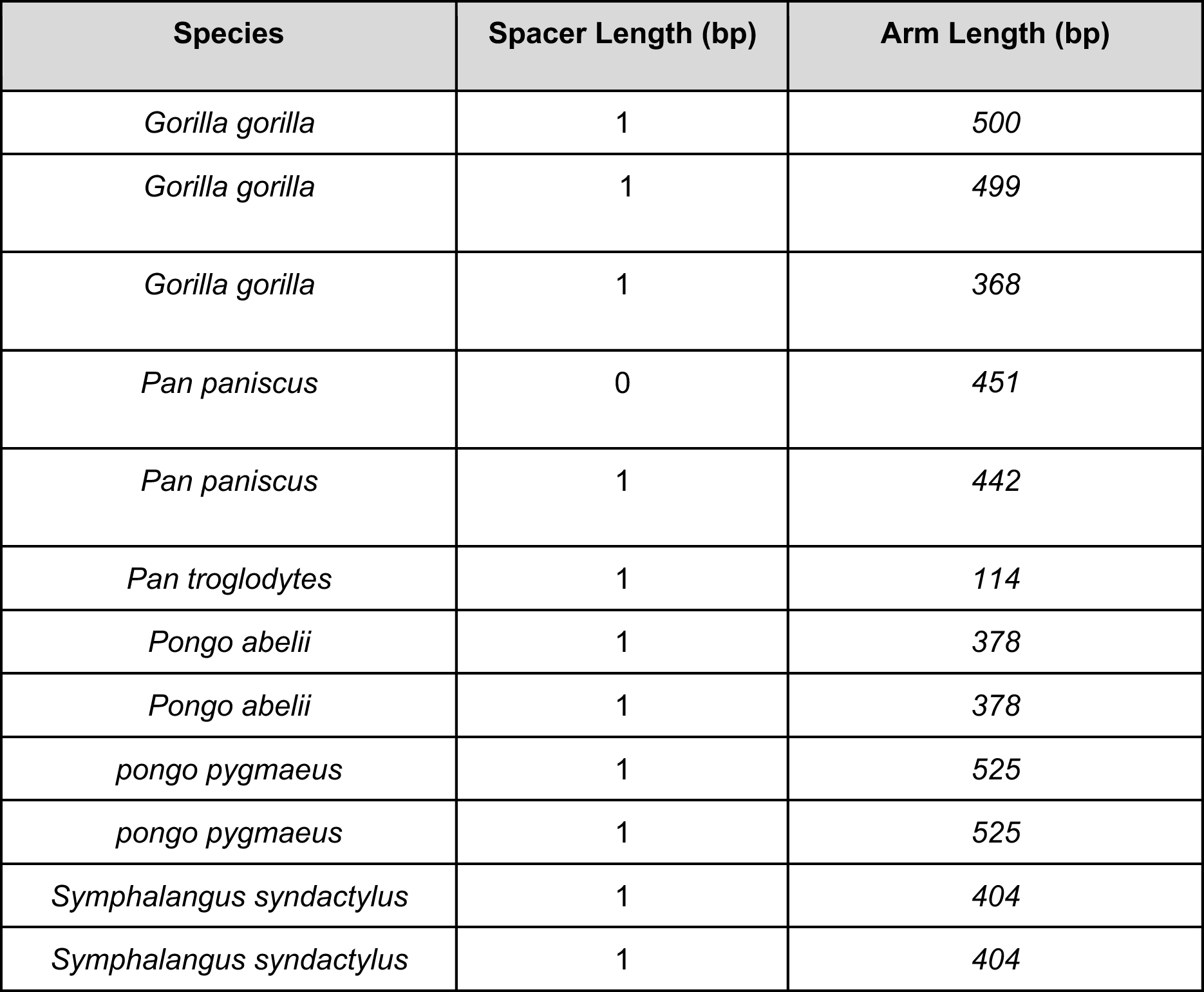
Longest H-DNA motifs across non-human great ape genomes.

| Species | Spacer Length (bp) | Arm Length (bp) |
| --- | --- | --- |
| *Gorilla gorilla* | 1 | 500 |
| *Gorilla gorilla* | 1 | 499 |
| *Gorilla gorilla* | 1 | 368 |
| *Pan paniscus* | 0 | 451 |
| *Pan paniscus* | 1 | 442 |
| *Pan troglodytes* | 1 | 114 |
| *Pongo abelii* | 1 | 378 |
| *Pongo abelii* | 1 | 378 |
| *pongo pygmaeus* | 1 | 525 |
| *pongo pygmaeus* | 1 | 525 |
| *Symphalangus syndactylus* | 1 | 404 |
| *Symphalangus syndactylus* | 1 | 404 |

